# Enzymatic Activity of PBP1B Compensates for β-lactam Mediated Inhibition of PBP2 in Cells Overproducing ppGpp

**DOI:** 10.1101/2025.06.12.659302

**Authors:** Sarah E. Anderson, Isabella E. Mack, Petra Anne Levin

**Affiliations:** Biology Department, William & Mary, Williamsburg, Virginia, USA; Department of Biology, Washington University in St. Louis, St. Louis, Missouri, USA

## Abstract

The alarmone (p)ppGpp (ppGpp) accumulates in response to starvation and other stress, leading to inhibition of multiple biosynthetic pathways and, at high concentrations, suppression of bacterial growth. Growth suppression by ppGpp is implicated in the formation of persister cells, which survive antibiotic challenge only to regrow once the drug is removed. However, there is also evidence that low levels of ppGpp contribute to resistance to certain cell wall-active antibiotics in actively growing cells. To characterize ppGpp’s contribution to antibiotic resistance, we measured MICs of a panel of β-lactams in actively growing *Escherichia coli* cells overexpressing a ppGpp synthase (*relA*\*). Cells engineered to modestly overproduce ppGpp exhibited up to 64-fold increases in resistance to PBP2-targeting β-lactams only, with mecillinam the most dramatically affected. Resistance required the transcription factor DksA and the class A penicillin binding protein (PBP) PBP1B. PBP1B variants defective for transpeptidase activity, glycosyltransferase activity, or both were incapable of mediating resistance, suggesting the full enzymatic activity of PBP1B is required for resistance. Transcriptomics revealed that ppGpp overproduction leads to increased expression of *lpoB*, which encodes an activator of PBP1B. LpoB was required for mecillinam resistance, with an *lpoB* deletion mutant exhibiting a loss of ppGpp-dependent resistance. Resistance was partially lost in an *lpoB* deletion strain expressing an LpoB-bypass variant of PBP1B (*mrcB*\*). Together these results support a model wherein overproduction of ppGpp favors cell wall synthesis by PBP1B, in part via increased expression of *lpoB*, allowing PBP1B to substitute for PBP2 when it is inhibited by β-lactams.

**Importance:** Antimicrobial resistance is an increasing global health threat, but the molecular mechanisms underlying resistance remain incompletely understood. This work clarifies the role of the alarmone ppGpp in mediating antibiotic resistance in the model bacterium *Escherichia coli*. Elevated levels of ppGpp were found to cause resistance to β-lactam antibiotics targeting the cell wall synthesis enzyme PBP2. Resistance required transcriptional regulation by ppGpp and enzymatic activity of the cell wall enzyme PBP1B. This work suggests that ppGpp causes resistance in part by increasing expression of *lpoB*, which encodes an activator of PBP1B. Because ppGpp levels are controlled by nutritional conditions, this work suggests that nutritional availability could impact the efficacy of antibiotics. Furthermore, this study adds to a number of studies demonstrating that cell wall synthesizing PBP enzymes can compensate for one another under certain conditions, leading to β-lactam resistance.

## Introduction

β-lactam antibiotics inhibit cell wall synthesis by inactivating penicillin binding proteins (PBPs). The model bacterium *Escherichia coli* encodes four major biosynthetic PBPs: the class A PBPs PBP1A (*mrcA*) and PBP1B (*mrcB*), which possess both glycosyltransferase and transpeptidase activity, as well as the essential class B transpeptidases PBP2 (*mrdA*) and PBP3 (*ftsI*) (1, 2). The class B enzymes are specialized for different modes of cell wall synthesis; PBP2 is a member of the elongasome that mediates cell elongation, while PBP3 is part of the divisome, required for division (3–5). β-lactams specific for PBP2 or PBP3 cause cell rounding or filamentation, respectively (5). The roles of the Class A PBPs are less clear; PBP1A interacts with both the elongasome and divisome, while PBP1B contributes to division and has been implicated in cell wall repair (6–11). Both enzymes require an outer membrane activator protein (LpoA for PBP1A and LpoB for PBP1B) for most of their functions (12, 13). Under standard laboratory conditions the class A PBPs are conditionally essential (14), indicating functional redundancy. However, PBP1A is required for maximal fitness in alkaline conditions and PBP1B becomes essential in acidic conditions (15), suggesting that these enzymes are also specialized for different environments. These enzymes can mediate environment- or mutation-dependent resistance to PBP2- and PBP3-targeting β-lactams (15–17), further supporting a model wherein functional redundancy among the PBPs allows them to substitute for one another in certain conditions.

The alarmones pppGpp and ppGpp [(p)ppGpp, hereafter referred to collectively as ppGpp] are also major drivers of environmental adaptation in bacteria. In *E. coli,* ppGpp levels are controlled by RelA (a monofunctional ppGpp synthetase) and SpoT (a bifunctional ppGpp synthetase/hydrolase) (18). ppGpp is produced at basal levels during balanced growth; under these conditions ppGpp is thought to contribute to general homeostatic control, including regulating the balance between longitudinal growth and division (19–21). ppGpp levels vary based on nutrient availability, with levels increasing during poor nutrient conditions. During starvation, ppGpp levels increase up to 100-fold, leading to a cessation of growth; this is known as the stringent response (SR) (19, 22–25).

In *E. coli* and its relatives, ppGpp acts via two regulatory pathways. ppGpp modulates transcription by directly binding to RNA polymerase (RNAP) and the RNAP-binding transcription factor DksA (26, 27), leading to differential expression of hundreds of genes during the SR (28). ppGpp also mediates post-translational regulation by directly binding at least fifty target proteins (25, 29, 30). Although ppGpp’s function in the SR is well understood, its contribution to survival in other stressful conditions, where its levels are lower and growth proceeds more or less normally, is less well characterized.

In addition to aiding survival during starvation, ppGpp also contributes to survival during exposure to antibiotics. Antibiotic tolerance, or the ability of bacteria to survive but not grow during antibiotic exposure, is associated with slowed growth rates (31, 32). ppGpp has been implicated in tolerance to multiple classes of antibiotics (32–36); this is thought to be due to ppGpp’s effects on growth rate, although the molecular mechanism is likely more nuanced (32, 36).

In contrast to tolerance, antibiotic resistance means that bacteria can both survive and grow in the presence of a drug. Mild elevations in ppGpp levels—low enough to reduce, but not completely inhibit, cell growth— are associated with resistance to mecillinam, a β-lactam that targets the essential, elongation specific class B PBP, PBP2 (37–43). Elevated ppGpp also causes resistance to multiple β-lactams in mutants overexpressing the L,D-transpeptidase gene *ldtD (ycbB)* (44). Overproduction of ppGpp alone does not increase resistance to ampicillin (which targets multiple PBPs), cephalexin (targets the essential, division-specific PBP3), or imipenem (targets multiple PBPs) (44–46), but the effect of ppGpp on resistance to other β-lactams is not known. Both ppGpp-mediated mecillinam resistance and broad spectrum β-lactam resistance in a ppGpp/LdtD overproducing strain depend on transcriptional regulation by ppGpp (47). Beyond this, the mechanism by which ppGpp causes mecillinam resistance is unclear.

In this work we systematically evaluated the effects of ppGpp on resistance to β-lactams targeting different PBPs. We found that ppGpp mediates resistance to PBP2-targeting β-lactams only, with mecillinam the most strongly affected. We discovered that ppGpp-mediated mecillinam resistance is dependent on both PBP1B and its activator LpoB. This finding adds to a growing number of examples of PBP1B-dependent mecillinam resistance (15, 17, 43), supporting a model wherein PBP1B can compensate for inhibition of PBP2 under certain conditions.

## Results

### Overproduction of ppGpp increases resistance to PBP2-targeting β-lactams

Previous work indicates that mild increases in ppGpp levels lead to mecillinam resistance in *E. coli* (37, 38, 41, 42); however, the effect of ppGpp on resistance to other β-lactams in the absence of mutations remains unclear. To determine whether ppGpp can cause resistance to β-lactams other than mecillinam, we quantified minimum inhibitory concentrations (MICs) for different β-lactams when ppGpp is overproduced. ppGpp overproduction was achieved using a plasmid (*prelA\**) encoding an IPTG-inducible copy of *relA\** (*relA1-455*), a truncated allele of *relA* encoding only the catalytic domain (48, 49). A plasmid encoding *relA’ (prelA’, relA1-331*), a further truncated and inactive allele of *relA*, was used as a negative control (50). Expression was induced with 10 µM IPTG, a concentration that caused only a modest ∼25% reduction in growth rate for cells harboring *prelA\** (**Supplemental Fig. S1A, C**).

We measured MICs of a panel of β-lactams that target different PBPs in *E. coli*. Similarly to published results, *prelA\** caused a 64-fold increase in the median mecillinam MIC (**Fig. 1A**). *prelA\** cells grew in the presence of 25 µg/ml mecillinam. For comparison, in previous work we found that the MIC for MG1655 was 0.4 µg/ml (15).

**Fig. 1.**
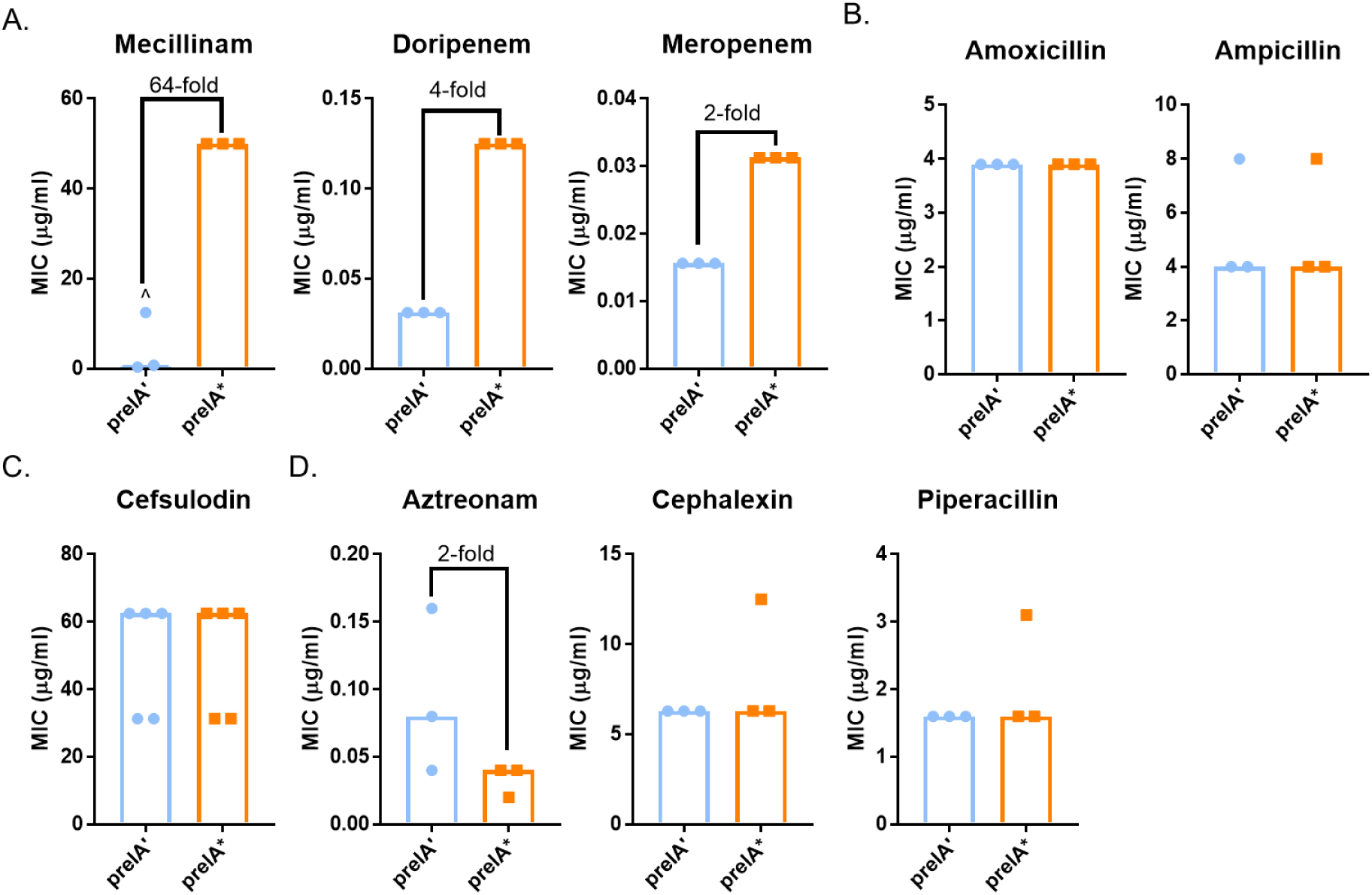
Overexpression of *relA\** increases resistance to PBP2-targeting β-lactams. **A.** Modest induction of *relA\** increases resistance to the PBP2-targeting β-lactams mecillinam, doripenem, and meropenem. **B-D.** Modest induction of *relA\** does not increase resistance to non-specific β-lactams (**B**), a PBP1A/PBP1B-targeting β-lactam (**C**), or PBP3-targeting β-lactams (**D**). Minimum inhibitory concentrations (MICs) for at least three biological replicates are shown along with median MICs and fold changes of median MICs. ^, growth skipped wells (see **Supplemental Fig. S2A**); the next concentration above the highest concentration of drug in which growth was observed was recorded as the MIC.

Intriguingly, the *prelA’* control exhibited an inconsistent response to mecillinam. While two biological replicates had MICs of 1.6 and 0.8 µg/ml respectively, one replicate grew at 0.4 µg/ml and at 1.6-6.3 µg/ml, but not 0.8µg/ml (**Supplemental Fig. S2**). We observed similar, but not identical, patterns of inconsistent growth in several subsequent experiments with mecillinam and once with doripenem (**Supplemental Fig. S2**). The recurring nature of this inconsistency indicates that it is unlikely to be due to pipetting error and may instead suggest heteroresistance or the accumulation of resistance-associated mutations. This phenomenon was considered beyond the scope of the current study and was not investigated further.

In *E. coli*, mecillinam specifically inhibits PBP2, which is also inactivated by meropenem and doripenem (45, 51). We observed modest but reproducible increases in the median MICs for doripenem (4-fold) and meropenem (2-fold) when *prelA\** was induced (**Fig. 1A**). We did not observe ppGpp-mediated increases in resistance to β-lactams targeting PBP1A and 1B, PBP3, or multiple PBPs (**Fig. 1B-1D**) (45, 51). Taken together, these results demonstrate that elevations in ppGpp levels increase resistance to PBP2-targeting β-lactams only, with mecillinam being most strongly affected.

To assess the contribution of basal levels of ppGpp to β-lactam susceptibility, we measured MICs for our panel of β-lactams against a strain unable to synthesize ppGpp (ppGpp^0^, Δ*relA spoT*::*cat*). ppGpp^0^ cells displayed modest ∼2-fold reductions in resistance to cefsulodin (targets PBP1A and 1B); doripenem, mecillinam, and meropenem (target PBP2); and piperacillin (targets PBP3). Resistance to aztreonam (targets PBP3) increased 2-fold, while resistance to the generalists amoxicilin and ampicillin, and the PBP3-targeting cephalexin, were unchanged (**Supplemental Fig. S3**). These values suggest that baseline levels of ppGpp are not a major contributor to intrinsic resistance under the conditions tested.

In addition to its reported effects on mecillinam resistance, ppGpp is also associated with β-lactam tolerance (52). To evaluate the impact of modest increases in ppGpp on tolerance, defined as the ability of bacterial cells to survive challenge with super-MIC concentrations of antibiotics, we measured survival of *prelA’* and *prelA\** cells in inhibitory concentrations of mecillinam, doripenem, cephalexin (PBP3-inhibitor), and ampicillin (non-specific). We found that *prelA\** reduced killing by all four antibiotics, although the differences for ampicillin were not statistically significant due to high levels of variability between replicates (**Supplemental Fig. S4**). This suggests that ppGpp-mediated β-lactam tolerance and resistance occur via different pathways, with resistance occurring to a much more limited subset of antibiotics.

### Resistance does not correlate with growth rate in mecillinam

It has been proposed that ppGpp causes β-lactam resistance by decreasing growth rate (43); however, it is unclear why a reduced growth rate would lead to resistance to PBP2-targeting β-lactams only. To determine if reduced growth rate in the presence of mecillinam correlates with resistance, we measured growth rates of *prelA’* and *prelA\** cells in the presence of a sub-inhibitory concentration of mecillinam (0.2 µg/ml). *prelA\** cells exhibited similar growth rates with and without mecillinam (1.80 ± 0.35 and 2.15 ± 0.35 doublings per hour, respectively) (**Supplemental Fig. S1**). In contrast, the growth rate of *prelA’* cells were reduced from 2.81 ± 0.02 doublings per hour without drug to 0.74 ± 0.18 doublings per hour with drug (**Supplemental Fig. S1**). This slowed growth rate may have been due to killing of some cells in the population, or to partial inhibition of PBP2 slowing elongation in all cells. As the more sensitive *prelA’* cells grew more slowly in sub-inhibitory concentrations of drug, this demonstrates that resistance does not correlate with reduced growth rate.

### ppGpp-mediated resistance requires DksA

ppGpp modulates global biosynthesis via transcriptional and post-translational pathways. Transcriptional regulation by ppGpp occurs in concert with the transcription factor DksA, which facilitates binding of ppGpp to RNAP (26).

Recent work suggests that ppGpp mediates mecillinam resistance through RNAP, as mutations in RNAP confer mecillinam resistance in the absence of ppGpp (47). To confirm whether ppGpp’s role in transcription is important for PBP2-targeting β-lactam resistance, we expressed *prelA’* and *prelA\** in a *dksA*::*kan* mutant and measured effects on MICs. The *dksA*::*kan* strain exhibited no ppGpp-dependent resistance to doripenem or meropenem, and showed only a two-fold increase in mecillinam resistance when *prelA\** was expressed (**Fig. 2**). This demonstrates that DksA is required for ppGpp to mediate β-lactam resistance, strongly suggesting that ppGpp mediates resistance through its effects on transcription.

**Figure 2.**
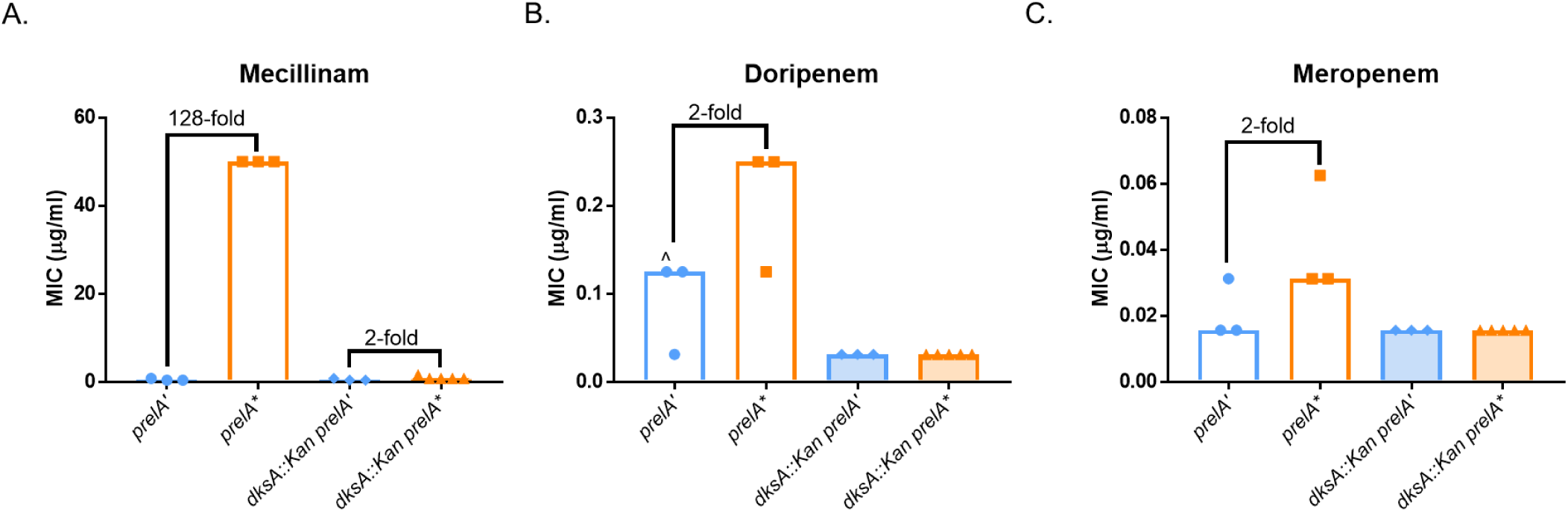
ppGpp-mediated resistance requires *dksA*, suggesting that ppGpp mediates resistance through transcription. Effects of deletion of *dksA* on ppGpp-dependent MICs of mecillinam (**A**), doripenem (**B**), and meropenem (**C**). MIC values for at least three replicates, median MICs, and fold changes of median MICs are shown. ^, growth skipped wells (see **Supplemental Fig. S2B**); the next concentration above the highest concentration of drug in which growth was observed was recorded as the MIC.

### ppGpp-mediated resistance depends on PBP1B

PBP1B also contributes to mecillinam resistance and tolerance (14–17). To determine whether PBP1B or the other major class A PBP, PBP1A, are involved in ppGpp-mediated resistance, we expressed *prelA’* and *prelA\** in Δ*mrcA* (PBP1A) and Δ*mrcB* (PBP1B) mutants and measured MICs of PBP2-targeting β-lactams. Deletion of *mrcB* fully eliminated ppGpp-mediated mecillinam and meropenem resistance, and decreased ppGpp-dependent doripenem resistance (**Fig. 3**). While *prelA\** caused a four-fold increase in doripenem resistance in the control strain, this was reduced to a two-fold increase in the Δ*mrcB* strain. Interestingly, the Δ*mrcB prelA’* and *prelA\** strains both exhibited lower doripenem MICs than their respective controls, suggesting that PBP1B also contributes to intrinsic doripenem resistance independently of ppGpp (**Fig. 3B**). Taken together, these results show that PBP1B contributes to ppGpp-mediated β-lactam resistance.

**Figure 3.**
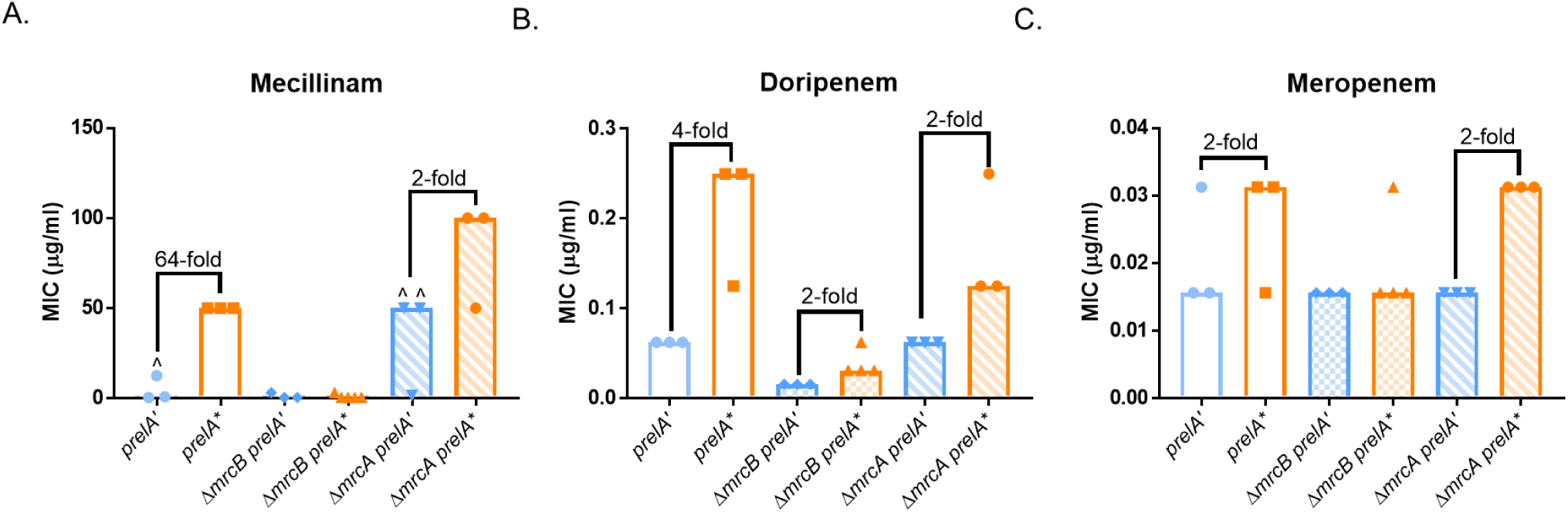
ppGpp-mediated resistance requires PBP1B. Effects of deletion of *mrcB* (encoding PBP1B) and *mrcA* (encoding PBP1A) on ppGpp-dependent MICs of mecillinam (**A**), doripenem (**B**), and meropenem (**C**). MICs values for at least three replicates, median MICs, and fold changes of median MICs are shown. ^, growth skipped wells (see **Supplemental Fig. S2C**); the next concentration above the highest concentration of drug in which growth was observed was recorded as the MIC.

In contrast, PBP1A does not appear to contribute to ppGpp-mediated resistance. The Δ*mrcA prelA\** strain was highly mecillinam resistant, with a median MIC of 100 µg/ml (**Fig. 3A**). We saw high rates of inconsistent growth in mecillinam for the Δ*mrcA prelA’* strain, which elevated the MIC for this strain to 50 µg/ml (**Fig. 3A, Supplemental Fig. S2C**). The Δ*mrcA prelA\** strain exhibited similar meropenem and doripenem MICs to the *prelA\** control (**Fig. 3B, C**). Overall, these results suggest that PBP1B, but not PBP1A, is required for ppGpp-mediated resistance to β-lactams, particularly mecillinam.

As a bifunctional class A PBP, PBP1B exhibits both transpeptidase (TPase) and glycosyltransferase (GTase) activity. To determine which enzymatic activities of PBP1B are required for resistance, we expressed *prelA*\* in Δ*mrcB* strains expressing separation-of-function alleles of *mrcB* from a plasmid. Mutant variants lacked TPase activity (*pmrcB-TP\**, *mrcB*S510A), GTase activity (*pmcrB-GT\**, *mrcB*E233Q) or both (*pmrcB-GT*TP\**, *mrcB*E233Q,S510A) (53). As expected, deletion of *mrcB* reduced *prelA\**-mediated mecillinam, doripenem, and meropenem resistance in a strain expressing a pBAD33 control plasmid (**Fig. 4**). Expression of wild-type *mrcB* from a plasmid (*pmcrB-WT*) fully complemented resistance to all three antibiotics.

**Figure 4.**
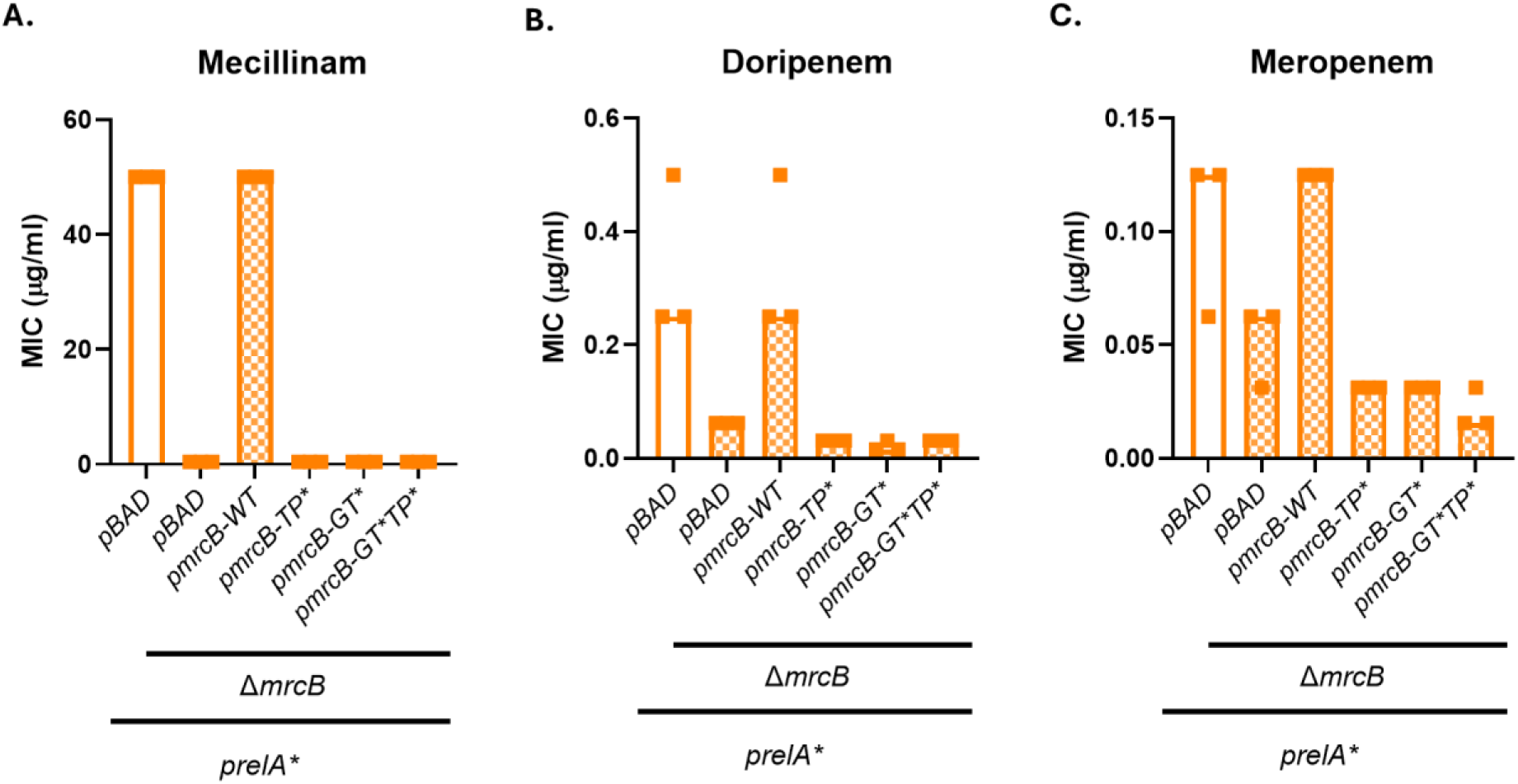
Separation-of-function *mrcB* mutants are incapable of supporting ppGpp-mediated resistance. Effects of disruption of PBP1B transpeptidase activity (TP*), glycotransferase activity (GT*), or both enzymatic activities (GT*TP*) on ppGpp-dependent MICs of mecillinam (A), doripenem (B), and meropenem (C). MIC values for at least three replicates and median MICs are shown.

However, the three PBP1B separation-of-function mutants were unable to restore the ppGpp-dependent resistance phenotype (**Fig. 4**). It should be noted that a disruption of GTase activity in PBP1B dramatically reduces TPase activity (54–56). Therefore, this result demonstrates that TPase enzymatic function of PBP1B is required for ppGpp-mediated antibiotic resistance, and suggests that GTase activity may also contribute.

### Overproduction of ppGpp causes mecillinam resistance in part by increasing *lpoB* expression

The simplest explanation for our data so far is that ppGpp mediates PBP1B activation through DksA-dependent transcription. To determine whether DksA and PBP1B affect resistance through the same pathway, we expressed *prelA’* and *prelA\** in a Δ*mrcB dksA*::*kan* double mutant and compared its resistance to each single mutant. The effects of *mrcB* and *dksA* on resistance were not additive in the double mutant (**Supplemental Fig. S5**), suggesting that *mrcB* and *dksA* impact ppGpp-mediated resistance through the same pathway. This result supports a model wherein ppGpp impacts resistance through a transcriptional pathway involving PBP1B.

To uncover the transcriptional pathway by which ppGpp contributes to β-lactam resistance, we performed RNA-sequencing with *prelA’* and *prelA\** strains induced with 10 µM IPTG (**Supplemental Data File 1**). Although many published studies have looked at gene expression changes due to the stringent response, we wanted to determine the effect of chronic, mild overproduction of ppGpp on gene expression. We found over 1,000 differentially regulated genes between *prelA\** and *prelA’* (**Supplemental Data File 1**). While we did not find changes in gene expression of *mrcB*, we did observe a statistically significant, ∼2-fold increase in expression of *lpoB*, which encodes the PBP1B activator protein (12, 13) (**Supplemental Data File 1**). Increased *lpoB* expression has also been reported following acute, high-level overproduction of ppGpp (28). A two-fold increase in *lpoB* expression has previously been reported to result in a 192-fold increase in mecillinam resistance (17). This raises the possibility that ppGpp may contribute to β-lactam resistance by modestly increasing expression of *lpoB*.

To assess whether LpoB is required for ppGpp-mediated resistance, we measured the effect of *prelA\** in an *lpoB*::*kan* mutant. This mutant exhibited only a two-fold increase in the median mecillinam MIC when *prelA\** was induced, demonstrating that LpoB is required for ppGpp-mediated mecillinam resistance (**Fig. 5A**). Our results for doripenem and meropenem were less clear. The *lpoB*::*kan* mutation reduced doripenem resistance in cells expressing *prelA’* and *prelA\**, although *prelA\** still caused a two-fold increase in resistance in this mutant background (**Fig. 5B**). Deletion of *lpoB* did not affect the modest two-fold increase in meropenem resistance caused by *prelA\** (**Fig. 5C**). These results demonstrate that LpoB is required for ppGpp-dependent mecillinam resistance.

**Figure 5.**
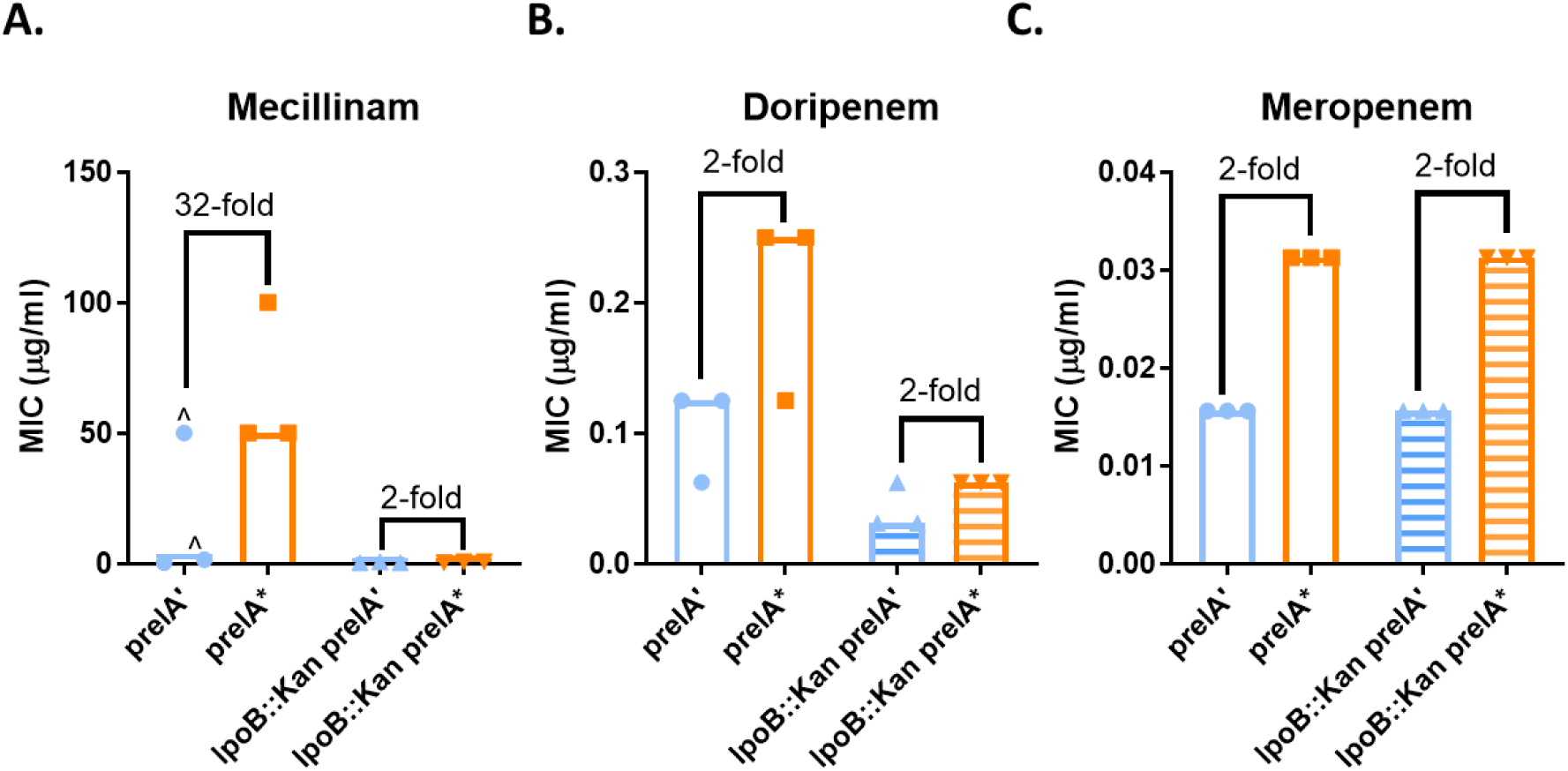
*lpoB* is required for ppGpp-mediated mecillinam resistance. Effects of an *lpoB*::*kan* deletion on ppGpp-mediated resistance to mecillinam (**A**), doripenem (**B**), and meropenem (**C**). MIC values for at least three replicates, median MICs, and fold changes of median MICs are shown. ^, growth skipped wells (see **Supplemental Fig. S2D**); the next concentration above the highest concentration of drug in which growth was observed was recorded as the MIC.

As LpoB is typically required for enzymatic activity of PBP1B, it is unclear whether the effect of the *lpoB*::*kan* mutation on mecillinam resistance is due to a loss of ppGpp signaling to PBP1B through LpoB, or to a general loss of PBP1B activity. To differentiate between these two possibilities, we took advantage of an LpoB-bypass mutant of PBP1B (*mrcBE313D*, designated *mrcB\**) (57). This mutation mimics the effect of LpoB activation on PBP1B, although PBP1B* appears to still be capable of interacting with LpoB (57, 58). We hypothesized that if ppGpp is causing β-lactam resistance by increasing *lpoB* expression, then an *mrcB\** strain should exhibit ppGpp-dependent resistance, while a *mrcB* lpoB*::*kan* mutant should not.

Our results suggest that ppGpp-dependent mecillinam resistance is partially dependent on signaling through LpoB. The *mrcB\** mutant exhibited a 25-fold increase in median mecillinam resistance due to *prelA\**, this was reduced to a 3-fold increase in the *mrcB* lpoB*::*kan* mutant (**Fig. 6A**). However, the *mrcB* lpoB*::*kan prelA’* control strain exhibited high rates of inconsistent growth at higher mecillinam concentrations (**Supplemental Fig. S2E**), inflating the MICs for this strain and contributing to the small fold change due to *prelA\**. Even so, the median mecillinam MIC for the *mrcB* lpoB*::*kan prelA\** strain was 5-fold lower than that for the *mrcB* prelA\** strain. This reduction in MIC suggests that LpoB does contribute to mecillinam resistance even in the *mrcB\** strain. However, the fact that the *mrcB* lpoB*::*kan* strain still responds to mecillinam suggests that additional, unidentified mechanism(s) also contribute to ppGpp- and PBP1B-dependent mecillinam resistance.

**Figure 6.**
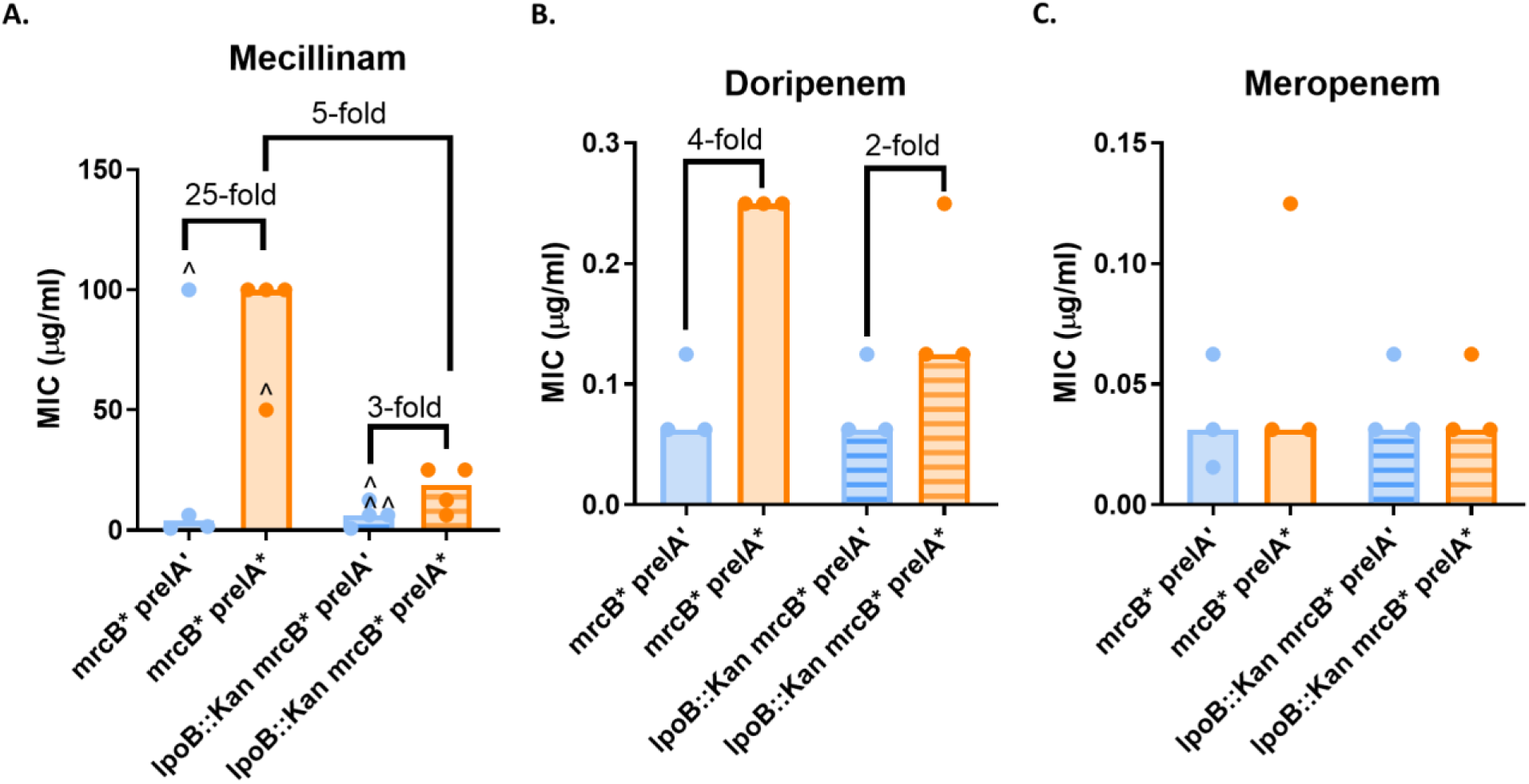
ppGpp mediates mecillinam resistance in part via LpoB. Effect of an *lpoB*::*kan* mutation on ppGpp-mediated resistance in a strain expressing an LpoB-bypass *mrcB\** mutation. Resistance was measured to mecillinam (**A**), doripenem (**B**), and meropenem (**C**). MIC values for at least three independent replicates, median MICs, and fold changes of median MICs are shown. ^, growth skipped wells (see **Supplemental Fig. S2E**); the next concentration above the highest concentration of drug in which growth was observed was recorded as the MIC.

LpoB may also contribute to ppGpp-dependent doripenem resistance in the *mrcB\** strain. We found a 4-fold increase in median MIC due to *prelA\** expression in *mrcB\** cells; this was reduced to a 2-fold increase in the *lpoB*::*kan mrcB\** strain (**Fig. 6B**). Surprisingly, the *mrcB\** and *lpoB*::*kan mrcB\** strains both failed to exhibit a reproducible increase in meropenem resistance due to *prelA\** (**Fig. 6C**), so the contribution of *lpoB* to ppGpp- and PBP1B-dependent meropenem resistance could not be further explored.

## Discussion

This study demonstrates that ppGpp mediates resistance to a subset of β-lactam antibiotics via a growth-rate independent mechanism. We have confirmed previous findings that ppGpp is responsible for mecillinam resistance and established that ppGpp can cause a mild increase in resistance to doripenem and meropenem (**Fig. 1A**). These three antibiotics preferentially target PBP2 in *E. coli* (45, 51). This resistance requires the transcription factor DksA (**Fig. 2**) and catalytic activity by the class A PBP PBP1B (**Fig. 3**, **4**). Our results suggest that ppGpp mediates mecillinam resistance in part by upregulating expression of *lpoB* (**Fig. 5**, **6, Supplemental Data File S1**), likely favoring cell wall synthesis by PBP1B when PBP2 is inhibited.

This work adds to a growing body of evidence that PBP1B is able to compensate for the inhibition of PBP2 under certain conditions, facilitating growth under otherwise inhibitory concentrations of antibiotics. Previous work implicates overexpression of *lpoB* and *mrcB* as drivers of mecillinam resistance in *E. coli* clinical isolates harboring mutations in *cysB* (17), supporting PBP1B-mediated mecillinam resistance as a clinically relevant phenomenon. Loss of PBP1B enhances lysis by both mecillinam and A22, a drug inhibiting the elongasome protein MreB (16, 59, 60). Additionally, loss of PBP1B also enhances lysis in strains carrying temperature sensitive alleles of *mrdA* (PBP2) and other elongasome genes (61). PBP1B also mediates resistance to PBP2-targeting β-lactams in acidic growth media, although this phenotype also extends to PBP3-inhibiting drugs (see below) (15). Multiple studies indicate that cefsulodin (which inhibits both PBP1A and PBP1B) is synergistic with other β-lactams, including mecillinam (62–65), further supporting a protective role for PBP1B during PBP2 inhibition.

It is unclear how PBP1B is able to compensate for inhibition of PBP2, especially since PBP1B is involved in cell division and cell wall repair, while PBP2 is only required for elongation (3, 5, 6, 9–11). However, increased expression of divisome genes, especially *ftsZ*, can also increase mecillinam resistance and compensate for deletion of PBP2, suggesting that the proteins involved in division may be able to substitute for the elongasome in some circumstances (39, 66). It is also possible that PBP1B mediates resistance by increasing rates of cell wall repair (6). More research is required to distinguish between these two roles in the case of mecillinam resistance.

It is also not apparent why ppGpp does not mediate resistance to β-lactams that are not specific to PBP2. ppGpp overproduction can lead to broad-spectrum β-lactam resistance if the L,D-transpeptidase LdtD (YcbB) is simultaneously overexpressed (44, 47). This broad-spectrum resistance requires glycosyltransferase activity from PBP1B, but not transpeptidase activity (44). This is in contrast to the resistance to PBP2-targeting antibiotics observed here, which requires transpeptidase activity of PBP1B (**Fig. 5**). These findings demonstrate that during ppGpp overexpression, PBP1B is able to compensate for the loss of transpeptidase activity from PBP2, but upregulation of an additional transpeptidase (i.e. LdtD) is required to compensate for inhibition of other PBPs. This conclusion seemingly contradicts the observation that PBP1B is capable of mediating resistance to both PBP2- and PBP3-targeting antibiotics in low pH conditions (15). It is possible that PBP1B is more highly activated by low pH than by ppGpp overexpression, or that additional changes in the cell wall synthesis machinery at low pH underlie this discrepancy.

ppGpp overproduction causes a 64-fold increase in mecillinam resistance, while doripenem and meropenem resistance increase only 2-4-fold (**Fig. 1A**). The low level of doripenem and meropenem resistance due to ppGpp overproduction made it difficult to assess the contribution of LpoB to resistance to these drugs (**Fig. 5**, **6**). The differences between PBP2-targeting β-lactams may be due to the chemical classes of these drugs, with doripenem and meropenem belonging to the carbapenem class, while mecillinam is a penicillin. All three of these drugs share PBP2 as their specific PBP transpeptidase target (45). However, doripenem and meropenem have also been reported to bind PBP4 (both) and PBP7 (doripenem only) (45), class C PBPs that do not catalyze transpeptidation, and instead modify peptidoglycan in other ways (67). More work is needed to understand why the PBP2-targeting carbapenems are less responsive to overproduction of ppGpp compared with mecillinam.

Our results suggest that ppGpp mediates mecillinam resistance in part by increasing expression of *lpoB*. An LpoB-bypass variant of PBP1B exhibits reduced, but not eliminated, ppGpp-dependent resistance (**Fig. 6**), suggesting that ppGpp likely also contributes to resistance via additional pathway(s). Overproduction of ppGpp results in global transcriptional changes (**Supplemental Data File S1**), including changes in expression of many genes involved in cell wall synthesis, which may also be contributing to resistance. It is also likely that ppGpp influences resistance by indirectly activating FtsZ (20). ppGpp has been proposed to mediate β-lactam resistance by reducing expression of rRNA operons, although experimental downregulation of rRNA transcription was unable to recapitulate resistance without additional mutations in *rpoB* (47). As a pleiotropic regulator, it is unsurprising that ppGpp would affect the resistance phenotype through multiple mechanisms.

Finally, this work has implications for the use of mecillinam, both clinically and in research settings. While we are not aware of any clinical reports of mecillinam resistance arising solely from mutations that increase ppGpp levels, these mutations are easily isolated in a lab setting (37, 44, 68). Because ppGpp levels vary based on nutritional state, these results also suggest that nutrient poor environments could cause temporary phenotypic mecillinam resistance, without the accumulation of mutations. Indeed, reversible mecillinam resistance has been observed in wild-type *E. coli* starved for isoleucine and valine (40). Such transient resistance would not be easily detected in a clinical lab setting. Because ppGpp-mediated resistance is entirely dependent on PBP1B, our results suggest that cefsulodin administration may decrease the risk of ppGpp-mediated mecillinam resistance. Lastly, from a research perspective, mecillinam has previously been used as a model β-lactam for studying β-lactam mechanisms of action (69). The results shown here demonstrate that mecillinam is a very unusual β-lactam, at least from the perspective of ppGpp-mediated resistance, and may not be the best candidate as a model for this class of drug.

## Materials and Methods

### Bacterial strains, plasmids, and growth conditions

Bacterial strains, plasmids, and primers used in this study are detailed in **Supplemental Table S1**. All strains used were in the MG1655 background, which is referred to as “wild-type.” Selectable mutations were transferred between strains via P1 transduction, with transductants confirmed by PCR (for all mutants) and sequencing (for *mrcB\** only). Plasmids *prelA\** and *prelA’* were generated from *pALS10* and *pALS14*, respectively, by replacing the *bla* gene with a spectinomycin resistance gene from *pBS58* via *in vivo* assembly (IVA) cloning (50, 70, 71). Plasmids to overexpress separation-of-function *mrcB* alleles were generated by amplifying *mrcB* genes from the plasmids pUM1Bα* (TP*), pUM1BTG*α (GT*), pUM1BTG*α* (TP*GT*), and pUM1Bα (WT) and cloning them into the plasmid *pBAD33* with IVA cloning (53). Primers were obtained from Integrated DNA Technologies (Coralville, IA).

Experiments were performed in LB broth (1% tryptone, 1% NaCl, 0.5% yeast extract). *prelA\** and *prelA’* plasmids were induced by supplementation with 10 µM IPTG (Gold Bio, St. Louis, MO) and maintained with 100 µg/ml spectinomycin (Gold Bio or Millipore Sigma, St. Louis, MO). *pmrcB* plasmids were induced by addition of 0.2% arabinose (Millipore Sigma) and maintained with 30 µg/ml chloramphenicol (VWR, Radnor, PA). Selection for chromosomal mutations was performed using 50 µg/ml kanamycin (Gold Bio or Millipore Sigma) or 12.5 µg/ml tetracycline (Millipore Sigma).

### Minimum inhibitory concentrations

Antibiotics were obtained from Millipore Sigma or Cayman Chemical (Ann Arbor, MI) (doripenem). Cells were isolated from a single colony and grown in LB supplemented with IPTG, spectinomycin, arabinose, and/or chloramphenicol, if appropriate. Cultures were grown at 37°C with shaking until mid-exponential phase (OD600 = 0.2-0.6). A 96-well plate was prepared containing two-fold serial dilutions of antibiotics in LB supplemented with IPTG, spectinomycin, arabinose, and/or chloramphenicol, as necessary. Cells were inoculated into each well to a final OD600 of ∼0.0001. Plates were sealed with a Breathe-easy membrane (USA Scientific, Ocala, FL) and incubated at 37°C with shaking for 20 h. MICs were determined as the lowest contiguous concentration of drug that prevented visible bacterial growth.

### RNA-sequencing

Cultures were inoculated from a single colony and grown in LB supplemented with spectinomycin. Cultures were grown at 37°C with shaking until log phase and back-diluted to an OD600 of 0.01 in LB supplemented with spectinomycin and IPTG. Cultures were then grown to log phase and back-diluted again to 0.01 into the same conditions. Once cultures reached an OD600 of ∼0.2, samples were collected and mixed 8:1 with RNA stop solution (95% ethanol, 5% water-saturated phenol) on ice followed by centrifugation for 10 min at 4,000 rpm at 4°C. Pellets were flash frozen in a dry ice ethanol bath and stored at -80°C. RNA was prepared with the Trizol Plus RNA Purification Kit (Invitrogen, Waltham, MA) supplemented with Max Bacterial Enhancement Reagent (Invitrogen), according to the manufacturer’s instructions. Samples were treated with DNase I (Invitrogen) according to the manufacturer’s instructions. Illumina 12M RNA-sequencing with rRNA depletion and fold-change analysis was performed by SeqCenter (Pittsburgh, PA).

### Growth curves

Growth curves were performed identically to MICs with incubation for 20 h in an Epoch 2 plate reader. OD600 values were measured every 10 min. Doubling times were calculated from the exponential portion of each growth curve using the Doubling Time Cell Calculator ++ (https://doubling-time.com/compute_more.php). Growth rates (doublings per hour) were calculated by dividing 60/doubling time.

### Antibiotic killing curves

Cultures were inoculated from an isolated colony in LB supplemented with IPTG and spectinomycin. Cultures were grown at 37°C with shaking to mid-log phase (OD600 = 0.2-0.6) and diluted to an OD600 of 0.1. Antibiotic was added to cultures at the indicated concentration and cultures were incubated at 37°C with shaking. Serial dilutions of cultures were plated on LB supplemented with spectinomycin before antibiotic addition and after 1, 2, 3, 4, 5, 8, and 21 h following addition. CFUs were enumerated following at least 12 h incubation at 37°C, and percent survival was calculated at each time point compared to t = 0.

### ppGpp^0^ suppressor tests

ppGpp^0^ strains easily accumulate suppressor mutations in RNA polymerase genes (72). For all experiments using ppGpp^0^ strains, cultures were tested for the presence of suppressors at the same time that samples were collected for MICs. Suppressor tests were conducted as previously described (20). Data was only included from experiments where the frequency of suppressors was ≤ 10%.

### Quantification and statistical analyses

All experiments were performed with at least three biological replicates. Statistical tests were performed as indicated in figure legends using GraphPad Prism 7.

## Acknowledgements

We thank Dr. Thomas Bernhardt for his generous gift of the *mrcB*\* strain. We would like to thank members of the Levin and Anderson labs for their helpful feedback and discussion. This work was supported by NIH grants R35-GM127331 to P.A.L. and F32-GM143886 to S.E.A. S.E.A. was also supported by an FRC Faculty Research Grant from William & Mary.

---

**Supplemental Fig. S1.**
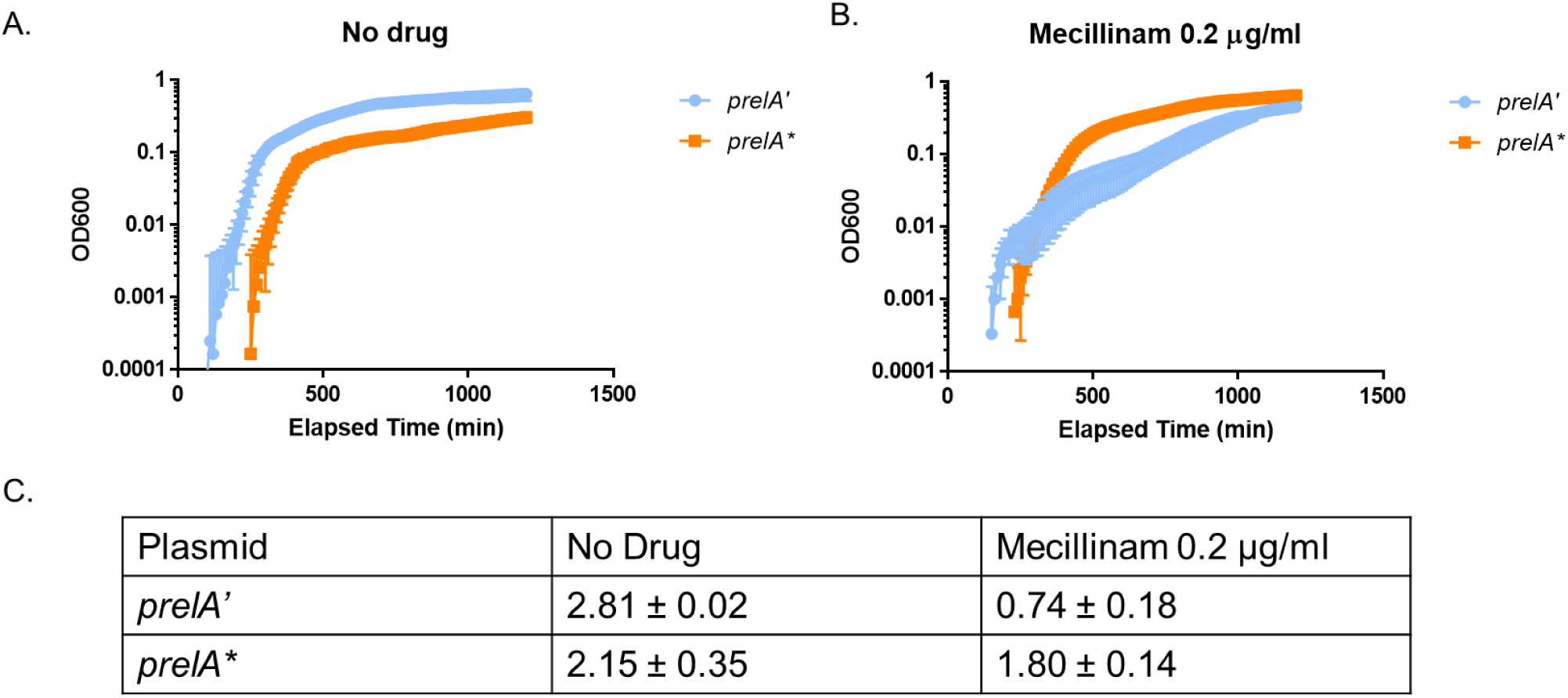
*prelA\** does not cause dramatic reductions in growth rate. **A.** Growth curves of *prelA’* and *prelA\** strains induced with 10 µM IPTG in the absence of drug. **B.** Growth curves of *prelA’* and *prelA\** strains induced with 10 µM IPTG in the presence of a sub- inhibitory concentration of mecillinam. **C**. Growth rates (doublings per hour) of *prelA’* and *prelA\** strains without drug and with mecillinam. Growth rates in sub-inhibitory concentrations of mecillinam do not correlate with minimum inhibitory concentrations of mecillinam (Fig. 1), suggesting that ppGpp does not cause resistance via lowering growth rate.

**Supplemental Figure S2.**
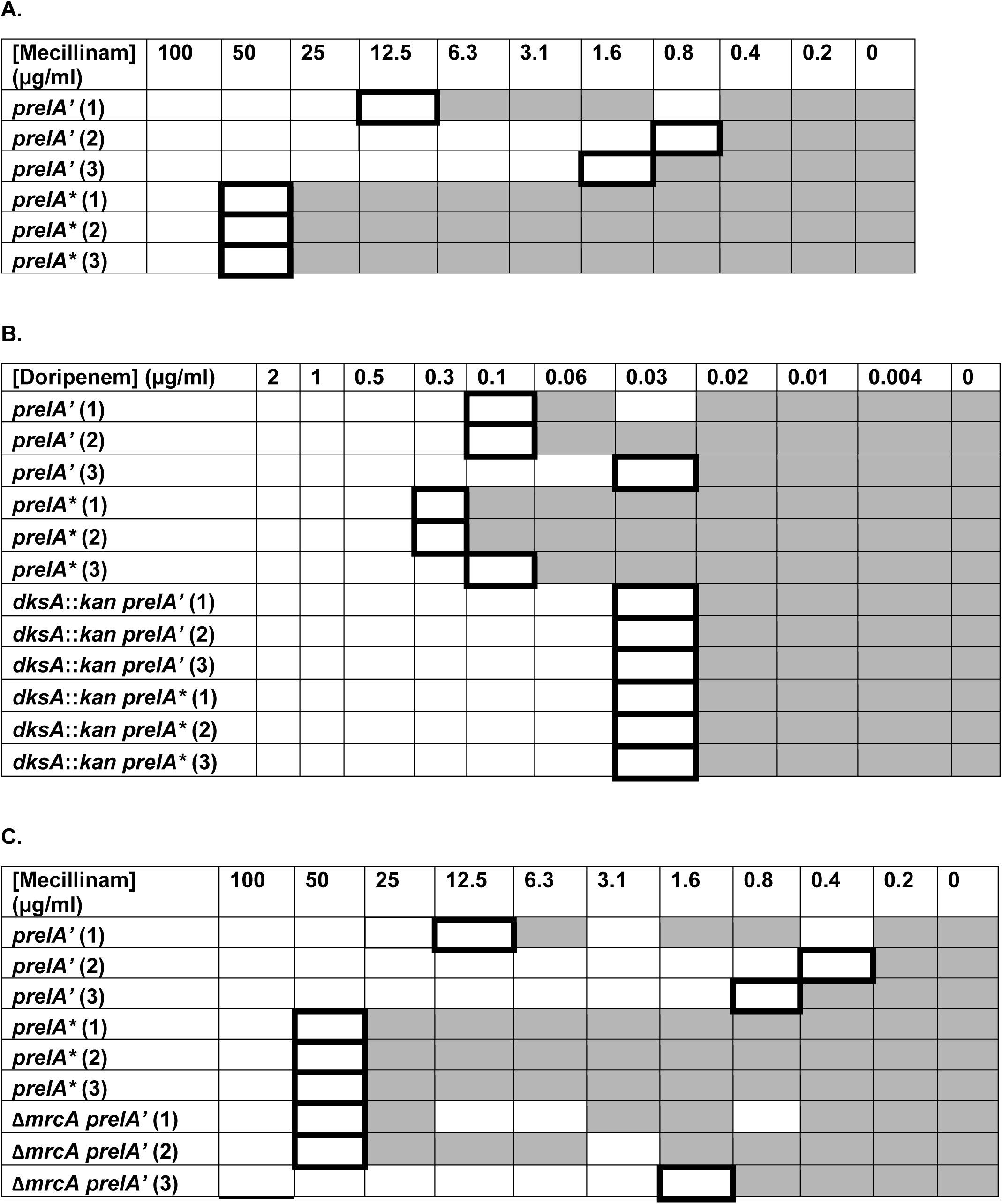
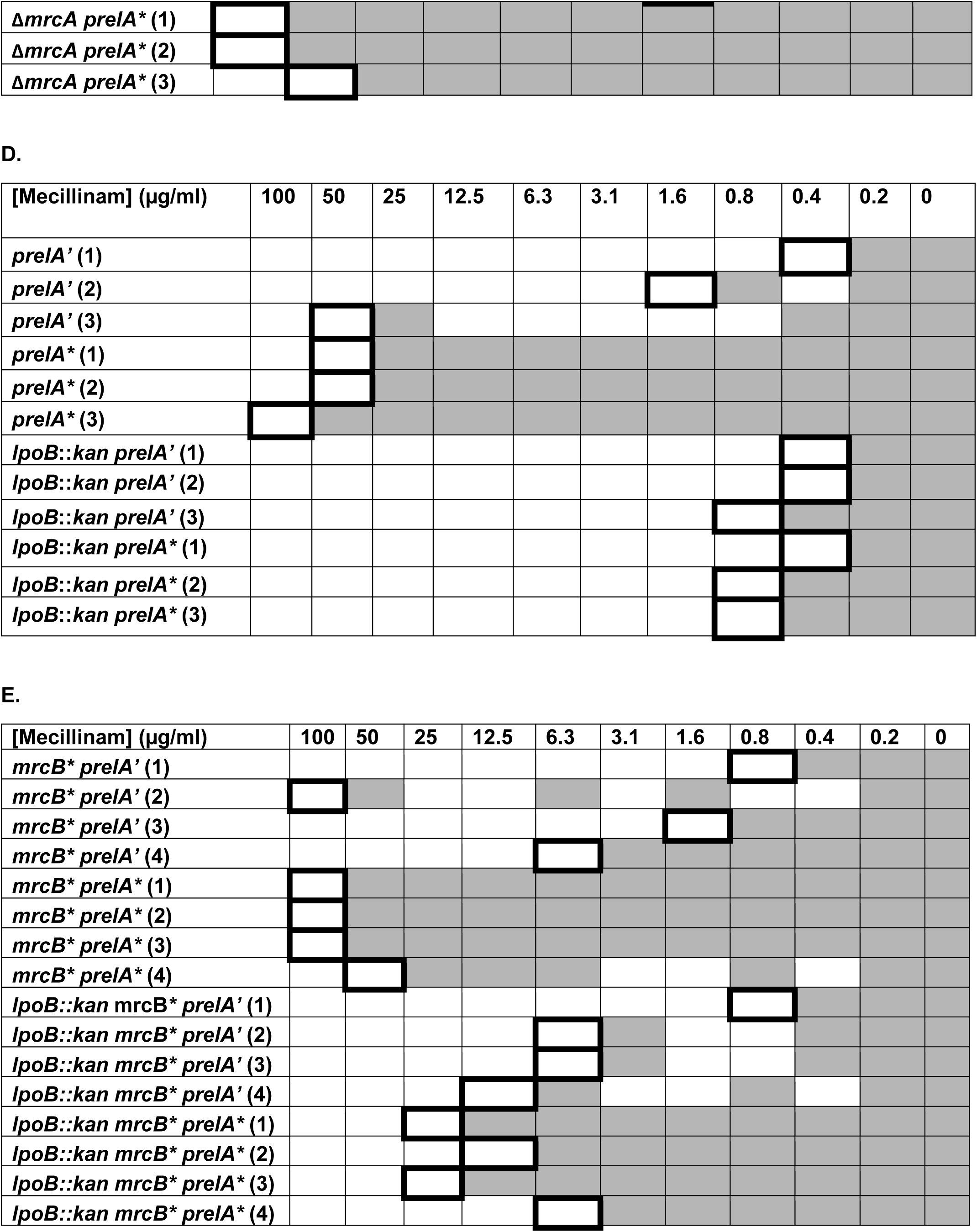
*prelA’* expressing cells exhibit occasional inconsistent growth and well skipping in mecillinam and doripenem MICs. Growth patterns for biological replicates used in **Fig. 1A** (**A**), **Fig. 2B** (**B**), **Fig. 3A** (**C**), **Fig. 5A** (**D**), **Fig. 6A** (**E**). Shading indicates visible growth at the indicated concentration of drug; bolded outlines indicate the concentration of drug that was recorded as the MIC.

**Supplemental Figure S3.**
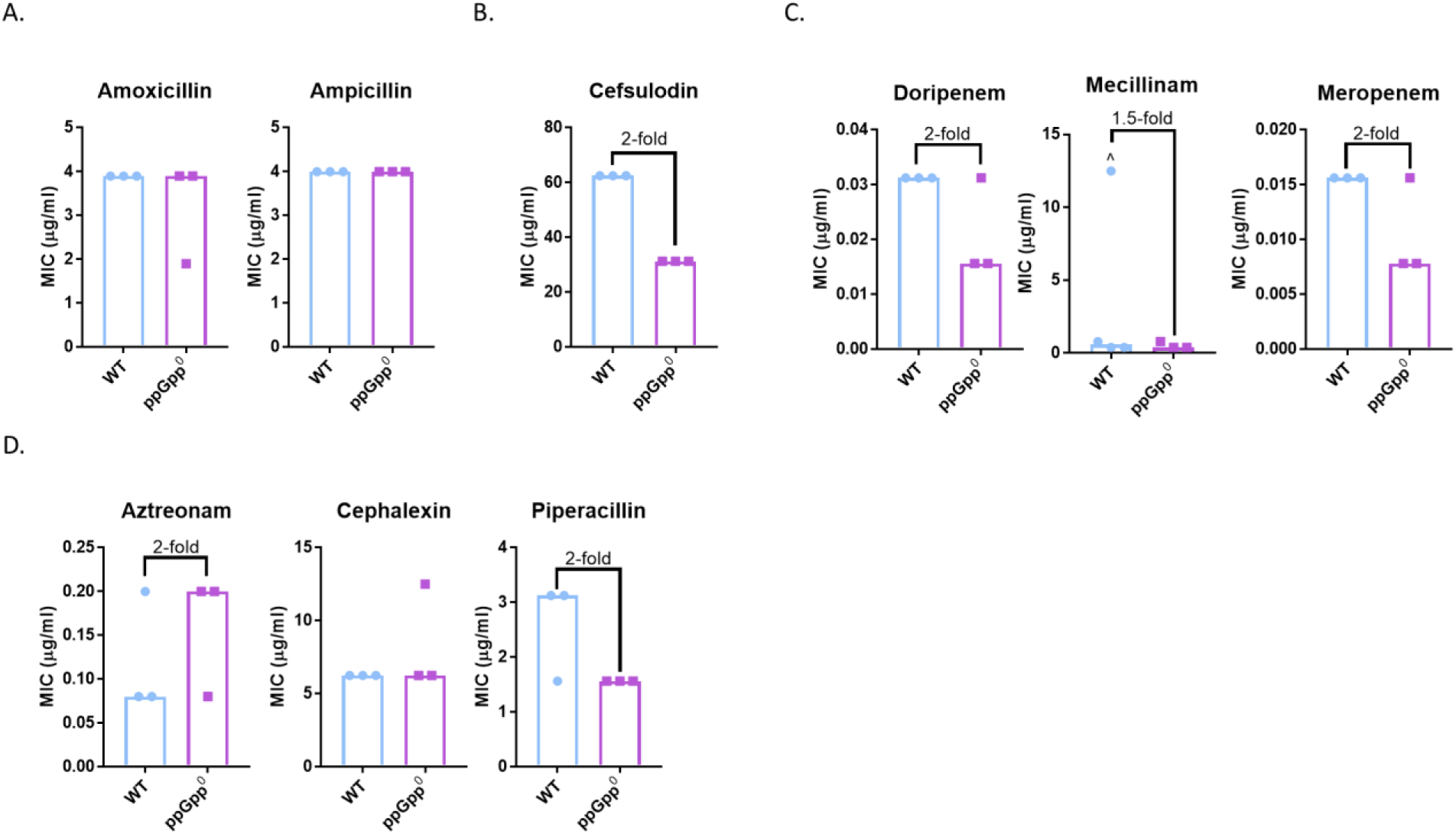
Loss of ppGpp (ppGpp^0^) does not greatly impact resistance to beta-lactams. MICs of ppGpp^0^ cells to β-lactams that are non-specific (**A**), target PBP1A/1B (**B**), target PBP2 (**C**), and target PBP3 (**D**). ^, growth skipped wells; the next concentration above the highest concentration of drug in which growth was observed was recorded as the MIC.

**Supplemental Figure S4.**
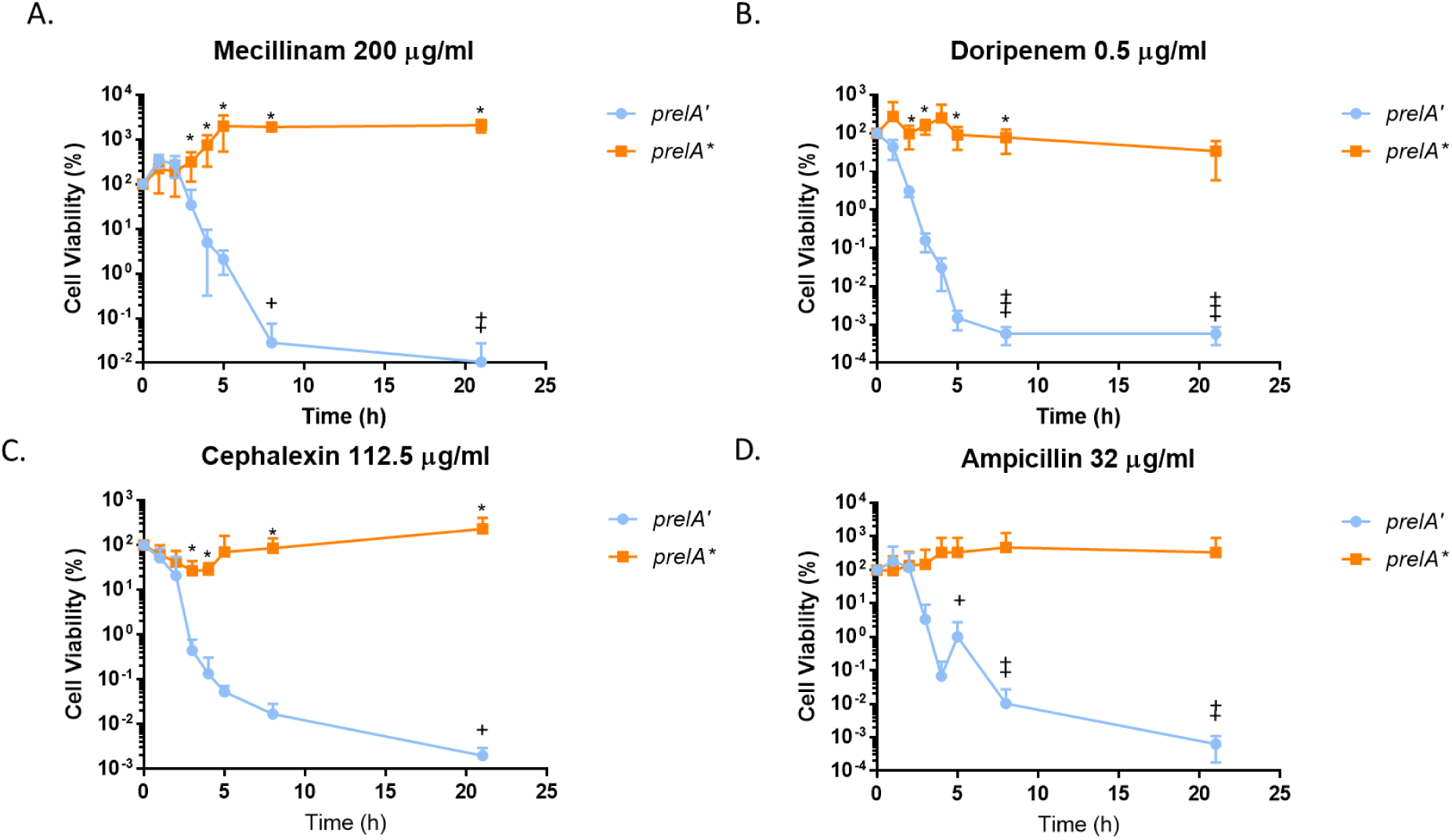
*prelA\** decreases killing by β-lactams with different targets. Survival of strains exposed to inhibitory concentrations of mecillinam (**A**), doripenem (**B**), cephalexin (**C**), and ampicillin (**D**). Data shown represent averages and standard deviations of three independent replicates. *, *p* ≤ 0.05 by two-tailed t-test; +, growth below limit of detection for one replicate; ‡, growth below limit of detection for two replicates; 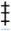, growth below limit of detection for three replicates. For samples below the limit of detection, the limit of detection was graphed.

**Supplemental Figure S5.**
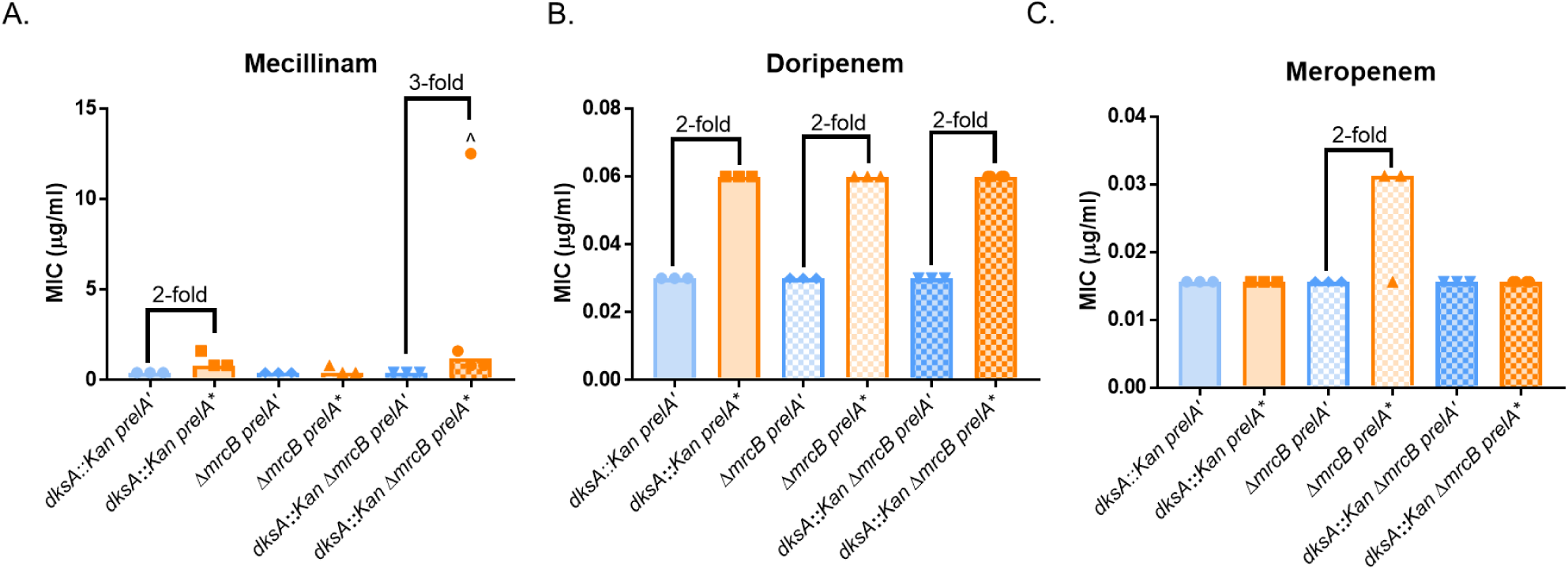
DksA and PBP1B contribute to ppGpp-mediated resistance through the same pathway. Effects of deletion of *dksA*, *mrcB*, and a double mutant on ppGpp- mediated resistance to mecillinam (**A**), doripenem (**B**), and meropenem (**C**). MIC values for at least three replicates, median MICs, and fold changes of median MICs are shown. ^, growth skipped wells; the next concentration above the highest concentration of drug in which growth was observed was recorded as the MIC.

## References

1. Sauvage E, Kerff F, Terrak M, Ayala JA, Charlier P. 2008. The penicillin-binding proteins: structure and role in peptidoglycan biosynthesis. FEMS microbiology reviews 32:234–258.

2. Pazos M, Peters K. 2019. Peptidoglycan. Sub-cellular biochemistry 92:127–168.

3. Cho H, Wivagg CN, Kapoor M, Barry Z, Rohs PDA, Suh H, Marto JA, Garner EC, Bernhardt TG. 2016. Bacterial cell wall biogenesis is mediated by SEDS and PBP polymerase families functioning semi-autonomously. Nature microbiology 1:16172.

4. Pogliano J, Pogliano K, Weiss DS, Losick R, Beckwith J. 1997. Inactivation of FtsI inhibits constriction of the FtsZ cytokinetic ring and delays the assembly of FtsZ rings at potential division sites. Proceedings of the National Academy of Sciences of the United States of America 94:559–64.

5. Spratt BG. 1975. Distinct penicillin binding proteins involved in the division, elongation, and shape of Escherichia coli K12. Proceedings of the National Academy of Sciences of the United States of America 72:2999–3003.

6. Vigouroux A, Cordier B, Aristov A, Alvarez L, Özbaykal G, Chaze T, Oldewurtel ER, Matondo M, Cava F, Bikard D, van Teeffelen S. 2020. Class-A penicillin binding proteins do not contribute to cell shape but repair cell-wall defects. eLife 9.

7. Banzhaf M, van den Berg van Saparoea B, Terrak M, Fraipont C, Egan A, Philippe J, Zapun A, Breukink E, Nguyen-Distèche M, den Blaauwen T, Vollmer W. 2012. Cooperativity of peptidoglycan synthases active in bacterial cell elongation. Molecular microbiology 85:179–94.

8. Pazos M, Peters K, Casanova M, Palacios P, VanNieuwenhze M, Breukink E, Vicente M, Vollmer W. 2018. Z-ring membrane anchors associate with cell wall synthases to initiate bacterial cell division. Nature Communications 9:5090.

9. Boes A, Kerff F, Herman R, Touze T, Breukink E, Terrak M. 2020. The bacterial cell division protein fragment (E)FtsN binds to and activates the major peptidoglycan synthase PBP1b. The Journal of biological chemistry 295:18256–18265.

10. Boes A, Olatunji S, Breukink E, Terrak M. 2019. Regulation of the Peptidoglycan Polymerase Activity of PBP1b by Antagonist Actions of the Core Divisome Proteins FtsBLQ and FtsN. mBio 10.

11. Navarro PP, Vettiger A, Hajdu R, Ananda VY, López-Tavares A, Schmid EW, Walter JC, Loose M, Chao LH, Bernhardt TG. 2025. The aPBP-type cell wall synthase PBP1b plays a specialized role in fortifying the Escherichia coli division site against osmotic rupture. bioRxiv 2025.04.02.646830.

12. Paradis-Bleau C, Markovski M, Uehara T, Lupoli TJ, Walker S, Kahne DE, Bernhardt TG. 2010. Lipoprotein Cofactors Located in the Outer Membrane Activate Bacterial Cell Wall Polymerases. Cell 143:1110–1120.

13. Typas A, Banzhaf M, van den Berg van Saparoea B, Verheul J, Biboy J, Nichols RJ, Zietek M, Beilharz K, Kannenberg K, von Rechenberg M, Breukink E, den Blaauwen T, Gross CA, Vollmer W. 2010. Regulation of peptidoglycan synthesis by outer-membrane proteins. Cell 143:1097–109.

14. Denome SA, Elf PK, Henderson TA, Nelson DE, Young KD. 1999. Escherichia coli mutants lacking all possible combinations of eight penicillin binding proteins: viability, characteristics, and implications for peptidoglycan synthesis. Journal of bacteriology 181:3981–93.

15. Mueller EA, Egan AJ, Breukink E, Vollmer W, Levin PA. 2019. Plasticity of Escherichia coli cell wall metabolism promotes fitness and antibiotic resistance across environmental conditions. eLife 8.

16. Grinnell A, Sloan R, Morgenstein RM. 2022. Cell density-dependent antibiotic tolerance to inhibition of the elongation machinery requires fully functional PBP1B. Communications biology 5:107.

17. Thulin E, Andersson DI. 2019. Upregulation of PBP1B and LpoB in cysB Mutants Confers Mecillinam (Amdinocillin) Resistance in Escherichia coli. Antimicrobial agents and chemotherapy 63.

18. Ronneau S, Hallez R. 2019. Make and break the alarmone: regulation of (p)ppGpp synthetase/hydrolase enzymes in bacteria. FEMS microbiology reviews 43:389–400.

19. Fernández-Coll L, Maciag-Dorszynska M, Tailor K, Vadia S, Levin PA, Szalewska-Palasz A, Cashel M. 2020. The Absence of (p)ppGpp Renders Initiation of Escherichia coli Chromosomal DNA Synthesis Independent of Growth Rates. mBio 11.

20. Anderson SE, Vadia SE, McKelvy J, Levin PA. 2023. The transcription factor DksA exerts opposing effects on cell division depending on the presence of ppGpp. mBio 14:e0242523.

21. Büke F, Grilli J, Cosentino Lagomarsino M, Bokinsky G, Tans SJ. 2021. ppGpp is a bacterial cell size regulator. Current biology : CB 10.1016/j.cub.2021.12.033.

22. Germain E, Guiraud P, Byrne D, Douzi B, Djendli M, Maisonneuve E. 2019. YtfK activates the stringent response by triggering the alarmone synthetase SpoT in Escherichia coli. Nature Communications 10:5763.

23. Dalebroux ZD, Swanson MS. 2012. ppGpp: magic beyond RNA polymerase. Nature reviews Microbiology 10:203–12.

24. Lee JW, Park YH, Seok YJ. 2018. Rsd balances (p)ppGpp level by stimulating the hydrolase activity of SpoT during carbon source downshift in Escherichia coli. Proceedings of the National Academy of Sciences of the United States of America 115:E6845–e6854.

25. Steinchen W, Zegarra V, Bange G. 2020. (p)ppGpp: Magic Modulators of Bacterial Physiology and Metabolism. Frontiers in microbiology 11:2072.

26. Ross W, Sanchez-Vazquez P, Chen AY, Lee JH, Burgos HL, Gourse RL. 2016. ppGpp Binding to a Site at the RNAP-DksA Interface Accounts for Its Dramatic Effects on Transcription Initiation during the Stringent Response. Molecular cell 62:811–823.

27. Ross W, Vrentas CE, Sanchez-Vazquez P, Gaal T, Gourse RL. 2013. The magic spot: a ppGpp binding site on E. coli RNA polymerase responsible for regulation of transcription initiation. Molecular cell 50:420–9.

28. Sanchez-Vazquez P, Dewey CN, Kitten N, Ross W, Gourse RL. 2019. Genome-wide effects on Escherichia coli transcription from ppGpp binding to its two sites on RNA polymerase. Proceedings of the National Academy of Sciences 116:8310–8319.

29. Wang B, Dai P, Ding D, Del Rosario A, Grant RA, Pentelute BL, Laub MT. 2019. Affinity-based capture and identification of protein effectors of the growth regulator ppGpp. Nature chemical biology 15:141–150.

30. Zhang Y, Zborníková E, Rejman D, Gerdes K. 2018. Novel (p)ppGpp Binding and Metabolizing Proteins of Escherichia coli. mBio 9.

31. Eng RH, Padberg FT, Smith SM, Tan EN, Cherubin CE. 1991. Bactericidal effects of antibiotics on slowly growing and nongrowing bacteria. Antimicrobial agents and chemotherapy 35:1824–1828.

32. Hobbs JK, Boraston AB. 2019. (p)ppGpp and the Stringent Response: An Emerging Threat to Antibiotic Therapy. ACS infectious diseases 5:1505–1517.

33. Westfall C, Flores-Mireles AL, Robinson JI, Lynch AJL, Hultgren S, Henderson JP, Levin PA. 2019. The Widely Used Antimicrobial Triclosan Induces High Levels of Antibiotic Tolerance In Vitro and Reduces Antibiotic Efficacy up to 100-Fold In Vivo. Antimicrobial agents and chemotherapy 63.

34. Harms A, Fino C, Sørensen Michael A, Semsey S, Gerdes K. 2017. Prophages and Growth Dynamics Confound Experimental Results with Antibiotic-Tolerant Persister Cells. mBio 8:10.1128/mbio.01964-17.

35. Germain E, Castro-Roa D, Zenkin N, Gerdes K. 2013. Molecular mechanism of bacterial persistence by HipA. Molecular cell 52:248–54.

36. Fung DK, Barra JT, Yang J, Schroeder JW, She F, Young M, Ying D, Stevenson DM, Amador-Noguez D, Wang JD. 2025. A shared alarmone-GTP switch controls persister formation in bacteria. Nat Microbiol 10.1038/s41564-025-02015-6.

37. Thulin E, Sundqvist M, Andersson DI. 2015. Amdinocillin (Mecillinam) resistance mutations in clinical isolates and laboratory-selected mutants of Escherichia coli. Antimicrobial agents and chemotherapy 59:1718–27.

38. Navarro F, Robin A, D’Ari R, Joseleau-Petit D. 1998. Analysis of the effect of ppGpp on the ftsQAZ operon in Escherichia coli. Molecular microbiology 29:815–23.

39. Vinella D, Joseleau-Petit D, Thévenet D, Bouloc P, D’Ari R. 1993. Penicillin-binding protein 2 inactivation in Escherichia coli results in cell division inhibition, which is relieved by FtsZ overexpression. Journal of bacteriology 175:6704–10.

40. Bouloc P, Vinella D, D’Ari R. 1992. Leucine and serine induce mecillinam resistance in Escherichia coli. Molecular & general genetics : MGG 235:242–6.

41. Joseleau-Petit D, Thévenet D, D’Ari R. 1994. ppGpp concentration, growth without PBP2 activity, and growth-rate control in Escherichia coli. Molecular microbiology 13:911–7.

42. Vinella D, D’Ari R, Jaffé A, Bouloc P. 1992. Penicillin binding protein 2 is dispensable in Escherichia coli when ppGpp synthesis is induced. The EMBO journal 11:1493–501.

43. Lai GC, Cho H, Bernhardt TG. 2017. The mecillinam resistome reveals a role for peptidoglycan endopeptidases in stimulating cell wall synthesis in Escherichia coli. PLoS genetics 13:e1006934.

44. Hugonnet JE, Mengin-Lecreulx D, Monton A, den Blaauwen T, Carbonnelle E, Veckerlé C, Brun YV, van Nieuwenhze M, Bouchier C, Tu K, Rice LB, Arthur M. 2016. Factors essential for L,D-transpeptidase-mediated peptidoglycan cross-linking and β-lactam resistance in Escherichia coli. eLife 5.

45. Kocaoglu O, Carlson EE. 2015. Profiling of β-Lactam Selectivity for Penicillin-Binding Proteins in <span class="named-content genus-species" id="named-content-1">Escherichia coli</span> Strain DC2. Antimicrobial agents and chemotherapy 59:2785–2790.

46. Hashizume T, Ishino F, Nakagawa J, Tamaki S, Matsuhashi M. 1984. Studies on the mechanism of action of imipenem (N-formimidoylthienamycin) in vitro: binding to the penicillin-binding proteins (PBPs) in Escherichia coli and Pseudomonas aeruginosa, and inhibition of enzyme activities due to the PBPs in E. coli. J Antibiot (Tokyo) 37:394–400.

47. Voedts H, Anoyatis-Pelé C, Langella O, Rusconi F, Hugonnet JE, Arthur M. 2024. (p)ppGpp modifies RNAP function to confer β-lactam resistance in a peptidoglycan-independent manner. Nature microbiology 9:647–656.

48. Schreiber G, Metzger S, Aizenman E, Roza S, Cashel M, Glaser G. 1991. Overexpression of the relA gene in Escherichia coli. The Journal of biological chemistry 266:3760–7.

49. Gropp M, Strausz Y, Gross M, Glaser G. 2001. Regulation of Escherichia coli RelA requires oligomerization of the C-terminal domain. J Bacteriol 183:570–579.

50. Svitil AL, Cashel M, Zyskind JW. 1993. Guanosine tetraphosphate inhibits protein synthesis in vivo. A possible protective mechanism for starvation stress in Escherichia coli. The Journal of biological chemistry 268:2307–11.

51. Shirley JD, Nauta KM, Carlson EE. 2022. Live-Cell Profiling of Penicillin-Binding Protein Inhibitors in Escherichia coli MG1655. ACS infectious diseases 8:1241–1252.

52. Dörr T. 2021. Understanding tolerance to cell wall-active antibiotics. Annals of the New York Academy of Sciences 1496:35–58.

53. Meisel U, Höltje JV, Vollmer W. 2003. Overproduction of inactive variants of the murein synthase PBP1B causes lysis in Escherichia coli. Journal of bacteriology 185:5342–8.

54. Egan AJF, Biboy J, van’t Veer I, Breukink E, Vollmer W. 2015. Activities and regulation of peptidoglycan synthases. Philosophical Transactions of the Royal Society B: Biological Sciences 370:20150031.

55. Terrak M, Ghosh TK, Van Heijenoort J, Van Beeumen J, Lampilas M, Aszodi J, Ayala JA, Ghuysen J-M, Nguyen-Distèche M. 1999. The catalytic, glycosyl transferase and acyl transferase modules of the cell wall peptidoglycan-polymerizing penicillin-binding protein 1b of Escherichia coli. Molecular Microbiology 34:350–364.

56. Bertsche U, Breukink E, Kast T, Vollmer W. 2005. *In Vitro* Murein (Peptidoglycan) Synthesis by Dimers of the Bifunctional Transglycosylase-Transpeptidase PBP1B from *Escherichia coli*\*. Journal of Biological Chemistry 280:38096–38101.

57. Markovski M, Bohrhunter JL, Lupoli TJ, Uehara T, Walker S, Kahne DE, Bernhardt TG. 2016. Cofactor bypass variants reveal a conformational control mechanism governing cell wall polymerase activity. Proceedings of the National Academy of Sciences of the United States of America 113:4788–93.

58. Ranjit DK, Jorgenson MA, Young KD. 2017. PBP1B Glycosyltransferase and Transpeptidase Activities Play Different Essential Roles during the De Novo Regeneration of Rod Morphology in Escherichia coli. Journal of bacteriology 199.

59. García del Portillo F, de Pedro MA. 1990. Differential effect of mutational impairment of penicillin- binding proteins 1A and 1B on Escherichia coli strains harboring thermosensitive mutations in the cell division genes ftsA, ftsQ, ftsZ, and pbpB. Journal of Bacteriology 172:5863–5870.

60. Park S, Cho H. 2022. The Tol-Pal System Plays an Important Role in Maintaining Cell Integrity During Elongation in Escherichia coli. Frontiers in microbiology 13:891926.

61. García del Portillo F, de Pedro MA. 1991. Penicillin-binding protein 2 is essential for the integrity of growing cells of Escherichia coli ponB strains. Journal of bacteriology 173:4530–2.

62. García del Portillo F, de Pedro MA, Joseleau-Petit D, D’Ari R. 1989. Lytic response of Escherichia coli cells to inhibitors of penicillin-binding proteins 1a and 1b as a timed event related to cell division. Journal of bacteriology 171:4217–21.

63. Laubacher ME, Ades SE. 2008. The Rcs phosphorelay is a cell envelope stress response activated by peptidoglycan stress and contributes to intrinsic antibiotic resistance. Journal of bacteriology 190:2065–74.

64. Powell JK, Young KD. 1991. Lysis of Escherichia coli by beta-lactams which bind penicillin-binding proteins 1a and 1b: inhibition by heat shock proteins. Journal of bacteriology 173:4021–6.

65. Sarkar SK, Dutta M, Kumar A, Mallik D, Ghosh AS. 2012. Sub-inhibitory cefsulodin sensitization of E. coli to β-lactams is mediated by PBP1b inhibition. PloS one 7:e48598.

66. Vinella D, Cashel M, D’Ari R. 2000. Selected amplification of the cell division genes ftsQ-ftsA-ftsZ in Escherichia coli. Genetics 156:1483–92.

67. van Heijenoort J. 2011. Peptidoglycan Hydrolases of Escherichia coli. Microbiology and Molecular Biology Reviews 75:636–663.

68. Bouloc P, Jaffé A, D’Ari R. 1988. Preliminary physiologic characterization and genetic analysis of a new Escherichia coli mutant, lov, resistant to mecillinam. Reviews of infectious diseases 10:905–10.

69. Cho H, Uehara T, Bernhardt TG. 2014. Beta-lactam antibiotics induce a lethal malfunctioning of the bacterial cell wall synthesis machinery. Cell 159:1300–11.

70. E Bi, J Lutkenhaus. 1990. FtsZ regulates frequency of cell division in Escherichia coli. Journal of bacteriology 172:2765–2768.

71. García-Nafría J, Watson JF, Greger IH. 2016. IVA cloning: A single-tube universal cloning system exploiting bacterial In Vivo Assembly. Scientific reports 6:27459.

72. Murphy H, Cashel M. 2003. Isolation of RNA Polymerase Suppressors of a (p)ppGpp Deficiency, p. 596–601. In Methods in Enzymology. Academic Press.

