## Supplemental Table 1 for "Enzymatic Activity of PBP1B Compensates for β-lactam Mediated Inhibition of PBP2 in Cells Overproducing ppGpp"

**Supplemental Table S1.** Bacterial strains, plasmids, and primers used in this study.

**Strains**

| **Designation** | **Genotype** | **Source** |
| --- | --- | --- |
| MG1655 (SEA1) | *rph1 ilvG rfb-50 λ- F-* | (1) |
| EAM696 (SEA1116) | MG1655 *mrcB*::*frt* | (2) |
| EAM899 (SEA1117) | MG1655 *mrcA*::*frt* | (2) |
| SEV161 (SEA1137) | MG1655 *dksA*::*kan* | (3) |
| CSW808 (SEA1094) | MG1655 *relA*::*frt* *spoT*::*cat* | (3) |
| SEA1011 | EAM696 *dksA*::*kan* | This study |
| EAM659 (SEA1072) | MG1655 *lpoB*::*kan* | (2) |
| MM43 | MG1655 *mrcB*(E313D) *yadC*::*Tn10* | (4) |
| SEA1069 | MG1655 (SEA1) *mrcB*(E313D) *yadC*::*Tn10* | This study |
| SEA1070 | EAM659 *mrcB*(E313D) *yadC*::*Tn10* | (4) This study |

**Plasmids**

| **Designation** | **Genotype** | **Source** |
| --- | --- | --- |
| *pALS13-spec* (*prelA**) | *lacI^q^* P*_tac_*-*relA*_1-455_ *spc^R^* | This study |
| *pALS14-spec* (*prelA’*) | *lacI^q^* P*_tac_*-*relA*_1-331_ *_­_spc^R^* | This study |
| *pBAD33* | *P_araBAD_ cm^R^* | (5) |
| *pmrcB-WT* | *P_araBAD_-mrcB cm^R^* | This study |
| *pmrcB-TP** | *P_araBAD_-mrcB_S510A_ cm^R^* | This study |
| *pmrcB-GT** | *P_araBAD_-mrcB_E233Q_ cm^R^* | This study |
| *pmrcB-GT*TP** | *P_araBAD_-mrcB_E233Q,S510A_ cm^R^* | This study |

**Primers**

| **Designation** | **Use** | **Sequence** | **Source** |
| --- | --- | --- | --- |
| oSEA211 | To generate *pALS13- and pALS14-spec* | ctgtcagaccaagtttactca | This study |
| oSEA212 |  | agagtttgtagaaacgcaaaa | This study |
| oSEA213 |  | aacttggtctgacaggttagacattatttgccgactac | This study |
| oSEA214 |  | cgtttctacaaactctggcttgttatgactgtttttttg | This study |
| oSEA240 | To generate *pmrcB* plasmids | GCTAGCCCAAAAAAACGGGTA | This study |
| oSEA241 |  | GAATTCGAGCTCGGTACCCG | This study |
| oSEA246 |  | TTTTTTTGGGCTAGCTTTCACACAGGAAACAGAATTC | This study |
| oSEA247 |  | ACCGAGCTCGAATTCCCCGCTTAGATGTTAATTACTACC | This study |

**References**

1.	Guyer MS, Reed RR, Steitz JA, Low KB. 1981. Identification of a sex-factor-affinity site in E. coli as gamma delta. Cold Spring Harbor symposia on quantitative biology 45 Pt 1:135–40.

2.	Mueller EA, Egan AJ, Breukink E, Vollmer W, Levin PA. 2019. Plasticity of Escherichia coli cell wall metabolism promotes fitness and antibiotic resistance across environmental conditions. eLife 8.

3.	Anderson SE, Vadia SE, McKelvy J, Levin PA. 2023. The transcription factor DksA exerts opposing effects on cell division depending on the presence of ppGpp. mBio 14:e0242523.

4.	Markovski M, Bohrhunter JL, Lupoli TJ, Uehara T, Walker S, Kahne DE, Bernhardt TG. 2016. Cofactor bypass variants reveal a conformational control mechanism governing cell wall polymerase activity. Proceedings of the National Academy of Sciences of the United States of America 113:4788–93.

5.	Guzman LM, Belin D, Carson MJ, Beckwith J. 1995. Tight regulation, modulation, and high-level expression by vectors containing the arabinose PBAD promoter. J Bacteriol 177:4121–4130.
